# Vascular injury associated with ethanol intake is driven by AT1 receptor and mitochondrial dysfunction

**DOI:** 10.1101/2023.09.05.556341

**Authors:** Wanessa M.C. Awata, Juliano V. Alves, Rafael M. Costa, Ariane Bruder-Nascimento, Shubhnita Singh, Gabriela S. Barbosa, Carlos Renato Tirapelli, Thiago Bruder-Nascimento

## Abstract

**Background:** Renin-angiotensin (Ang II)-aldosterone system (RAAS) is crucial for the cardiovascular risk associated with excessive ethanol consumption. Disturbs in mitochondria have been implicated in multiple cardiovascular diseases. However, if mitochondria dysfunction contributes to ethanol-induced vascular dysfunction is still unknown. We investigated whether ethanol leads to vascular dysfunction via RAAS activation, mitochondria dysfunction, and mitochondrial reactive oxygen species (mtROS).

**Methods:** Male C57/BL6J or mt-keima mice (6–8-weeks old) were treated with ethanol (20% vol./vol.) for 12 weeks with or without Losartan (10 mg/kg).

**Results:** Ethanol induced aortic hypercontractility in an endothelium-dependent manner. PGC1α (a marker of biogenesis), Mfn2, (an essential protein for mitochondria fusion), as well as Pink-1 and Parkin (markers of mitophagy), were reduced in aortas from ethanol-treated mice. Disturb in mitophagy flux was further confirmed in arteries from mt-keima mice. Additionally, ethanol increased mtROS and reduced SOD2 expression. Strikingly, losartan prevented vascular hypercontractility, mitochondrial dysfunction, mtROS, and restored SOD2 expression. Both MnTMPyP (SOD2 mimetic) and CCCP (a mitochondrial uncoupler) reverted ethanol-induced vascular dysfunction. Moreover, L-NAME (NOS inhibitor) and EUK 134 (superoxide dismutase/catalase mimetic) did not affect vascular response in ethanol group, suggesting that ethanol reduces aortic nitric oxide (NO) and H_2_O_2_ bioavailability. These responses were prevented by losartan.

**Conclusion:** AT_1_ receptor modulates ethanol-induced vascular hypercontractility by promoting mitochondrial dysfunction, mtROS, and reduction of NO and H_2_O_2_ bioavailability. Our findings shed a new light in our understanding of ethanol-induced vascular toxicity and open perspectives of new therapeutic approaches for patients with disorder associated with abusive ethanol drinking.

## 1. INTRODUCTION

Harmful ethanol consumption is a trigger for more than 200 disease and injury conditions, which result in 3 million deaths every year representing 5.3% of all deaths worldwide ^1^. During COVID-19 pandemic, the number of health problems related to ethanol consumption strikingly increased by 25% ^2^. Annual hospital costs in the United States of America (USA) for ethanol-dependent patients are approximately $249.0 billion resulting in a significant healthcare cost ^3^. We ^4–8^ and others ^9–12^ have demonstrated that excessive ethanol intake exacerbates the cardiovascular risk by promoting vascular dysfunction and inflammation, such abnormalities being dependent on a dysregulated redox signaling - exacerbated ROS formation, which involves a decompensated antioxidant response and/or exacerbated activity of oxidative enzymes ^6,13^.

The main sources of ROS in the vasculature are the enzyme β-nicotinamide adenine dinucleotide phosphate [NAD(P)H] oxidase (NOX), mitochondrial-derived superoxide (O_2_^•-^), uncoupled nitric oxide synthase (NOS), xanthine oxidase, cyclooxygenase, and myeloperoxidase ^14^. In the context of vascular injury associated with alcoholism, NOX enzymes-derived ROS are broadly studied and seem to be an attractive therapeutic target ^5^, while little is known on the participation of mitochondrial-derived ROS. Formation of mtROS is regulated by a fine arrangement between mitochondrial biogenesis, dynamic (fusion and fission), recycle (mitophagy), and oxidative phosphorylation (OXPHOS) ^15,16^. Ethanol induces mitochondrial dysregulation and/or mitochondrial-derived O_2_^•-^ in alveolar macrophage ^17,18^, heart, and T-cells ^19,20^. Although the connection between mitochondria dysfunction and mtROS generation has been described in non-vascular cells, whether ethanol consume induces mitochondria dysfunction and subsequently ROS formation and vascular dysfunction is still unknown.

The role of RAAS has been extensively studied on cardiovascular physiology and pathophysiology because of its important actions on water balance, and cardiorenal and vascular function, inflammation, and remodeling ^21^. We and others have previously demonstrated that overactivation of RAAS propagates vascular damage in rodents treated with ethanol by regulating ROS production and cyclooxygenase activation ^5,7,8,22–24^, whereas Ang II type 1 receptor (AT1R) blockade prevents chronic ethanol consumption-induced cardiomyopathy in dogs ^25^. Although the interface between chronic ethanol consumption, RAAS, and cardiovascular risk has been described, the downstream signaling pathways are still not fully elucidated.

Mitochondria quality is sensitive to Ang II effects in the context of endothelial dysfunction ^26,27^ and cardiac hypertrophy ^28^ and contractility ^29^, but whether the vascular changes associated with ethanol consumption is a RAAS and mitochondria crosstalk dependent manner is still unclear. Thus, we sought to investigate the consequences of ethanol consumption on mitochondrial quality of thoracic aortic and to analyze whether such changes may contribute to vascular injury. Therefore, we tested the hypothesis that ethanol consumption leads to a defective mitochondrion and mtROS overproduction, which in turn drive vascular dysfunction via an exacerbated RAAS activation. Of importance, understanding the vascular mechanisms underlying chronic ethanol consumption can help guide the development of novel cardiovascular therapeutic strategies for patients with disorder associated with abusive ethanol drinking.

## 2. METHODS

### 2.1. Ethical approval

Male isogenic mice of strains C57BL/6J (*Wild type*, WT) with 6 to 8 weeks (20 to 25 g) were purchased from the vivarium of the University of Pittsburgh were used. The mice were kept in the Rangos Research Building (room #4188), which has a temperature controlled by slipt-type air conditioning (22-24 ° C), automatic 12-hour light/dark cycle (lights on between 07:00 and 19:00 hours) with free access to water and food. The mice remained in a ventilated *rack* kept in groups of 4 animals per cage. All protocols were approved by the Institutional Animal Care and Use Committee (approval protocols numbers: 19065333 and 22061179) at University of Pittsburgh.

### 2.2. Experimental groups

Mice were chronically treated with 20% ethanol solution vol./vol. for 12 weeks, such treatment was based on our previous work, which describes a striking aortic oxidative stress ^30^.This treatment achieves a blood concentration (260 mg/dL) of ethanol in animals that resembles ethanol concentrations in heavy drinkers, which is sufficient to lead to cardiovascular changes^13,31–33^. The participation of RAAS in ethanol-induced vascular dysfunction was evaluated by treating mice with losartan (10 mg/kg, incorporated in food), a widely used selective AT_1_ receptor antagonist ^34^.

The animals were randomly distributed in 4 groups: 1) Control: animals had free access to water and received normal diet [losartan vehicle]; 2) Ethanol: animals had free access to a 20% ethanol solution (vol./vol.) and received normal diet [losartan vehicle]; 3) Losartan: animals had free access to water and received a normal diet_losartan (10 mg/kg) 4) Ethanol-losartan: animals had free access to a 20% ethanol solution (vol./vol.) and received a normal diet_losartan (10 mg/kg/day). The mice in the ethanol groups were conditioned to an adaptation period of 2 weeks in which they had free access exclusively to ethanol solutions of 5 and 10% (vol./vol.) in the first and second week, respectively. After adaptation, the mice had access exclusively to the 20% ethanol solution (vol./vol.) for 10 weeks ^35^. Body weight, food and water intake were measured twice a week. At the end of the 12th week, the animals were killed by CO_2_ saturation followed by rupture of the diaphragm, then heart, liver, and kidneys were isolated, weighed, and normalized by final body weight.

### 2.3. Design of losartan enriched diet

Since ethanol affects the food behavior, two different diets contained losartan were customized based on daily food intake (Dyets, Inc. PA-USA). Diet with losartan for control mice contained 0.058g/Kg of diet, while diet with losartan for mice treated with ethanol contained 0.076g/Kg. The amounts of losartan were calculated by a pilot study during which we observed that control mouse consumed approximately 4.2g/day and ethanol-treated mouse 3.5g/day.

### 2.4. Assessment of vascular function

After 12 weeks of treatment, thoracic aorta was harvested to perform studies of vascular reactivity. Vascular function was studied in rings (2mm) of thoracic aorta that were mounted in a wire myopgraph (DanyshMyoTechnology) to measure isometric tension recordings with PowerLab software (AD Instruments), as previously described^36^. Since ethanol can affect the function of endothelial cells, vascular smooth muscle cells, and perivascular adipose tissue (PVAT), we investigated the vascular contractility in aortic rings with intact [Endo (+)] and denuded endothelium [Endo (-)] with or without PVAT. The rings were placed in tissue baths containing Krebs-Henseleit solution (composition in mmol/L: 130 NaCl, 14.9 NaHCO_3_, 4.7 KCl, 1.18 KH_2_PO_4_, 1.17 MgSO_4_·, 5.5 glucose, 1.56 CaCl_2_ and 0.026 EDTA) gassed with 5% CO_2_: 95% O_2_, at 37°C and pH of 7.4. Cumulative concentration-response curves for phenylephrine (10^-10^ to 10^-4^ mol/L) in aorta and aorta+PVAT with or without endothelium were performed. To evaluate the mechanisms whereby ethanol consumption induces vascular disfunction and the participation of Ang II in these responses, concentration-response curves for phenylephrine were obtained after incubations for 30 min with manganese(III) tetrakis(1-methyl-4-pyridyl)porphyrin (MnTMPyP, 3 x 10 ^-2^ mmol/L, mitochondrial antioxidant – SOD-Mn mimetic), carbonyl cyanide m-chlorophenyl hydrazine (CCCP, 10^-3^ mmol/L, mitochondrial uncoupler), ethylbisiminomethylguaiacol manganese chloride (EUK-134, 10^-2^ mmol/L, synthetic superoxide dismutase and catalase mimetic) or NG-nitro-l-arginine methyl ester (L-NAME, 10^-1^ mmol/L, a nonselective NO synthase inhibitor). The choice of concentration of the inhibitors was chosen on previous reports ^37–40^. The maximal effect (Emax) and pD_2_ generated by phenylephrine was calculated from concentration–response curves, which were fitted using a non-linear interactive fitting program (Graph Pad Prism; GraphPad Software Inc., San Diego, CA, USA). The Emax correspond to the maximum effect elicited by phenylephrine and pD_2_ is the negative logarithm of the molar concentration of phenylephrine that promoted 50% of the maximum effect (EC_50_).

### 2.5. Histology

Thoracic aortas were collected and then fixed for 24 hours in 4% paraformaldehyde. The following step involved dehydration in 70% ethanol until the day of preparing the samples for histology. Aortas were embedded in paraffin and cut with a microtome and stained with hematoxylin and eosin (H&E). For image analysis, the thickness of the tunica media was measured (µm).

### 2.6. Quantitative real-time RT-PCR

RNeasy Mini Kit (Quiagen, Germantown, MD – USA) was used to extract mRNA from thoracic aorta. Complementary DNA (cDNA) was generated by reverse transcription polymerase chain reaction (RT-PCR) with SuperScript III (Thermo Fisher Waltham, MA USA). Reverse transcription was performed at 58°C for 50 min; the enzyme was heat inactivated at 85°C for 5 min, and real-time quantitative RT-PCR was performed with the PowerTrack™ SYBR Green Master Mix (Thermo Fisher, Waltham, MA USA). Sequences of genes as listed in supplementary table 1. Experiments were performed in a QuantStudio™ 5 Real-Time PCR System, 384-well (Thermo Fisher, Waltham, MA USA). Data were quantified by 2ΔΔ Ct and are presented by fold changes.

### 2.7. Protein expression evaluation by Western Immunoblotting

Thoracic aortas were frozen in liquid nitrogen and homogenized in an assay buffer (RIPA) (30mmol/L HEPES, pH 7.4, 150mM NaCl, 1% Nonidet P-40, 0.5% sodium deoxycholate, 0.1% sodium dodecyl sulfate, 5mM EDTA, 1mM NaV0_4_, 50mM NaF, 1mM PMSF, 10% pepstatin A, 10 μg/mL leupeptin and 10 μg/mL aprotinin). Polyacrylamide gradient gel (BioRad Hercules, California – USA) was used for protein separation by electrophoresis and transferred to Immobilon-P poly (vinylidene fluoride) membranes. Blockade of non-specific binding sites was performed with 5% skim milk in tris-buffered saline solution with tween 20 (TBS-T) for 1h at 24°C. The membranes were incubated with specific antibodies overnight at 4 °C. The antibodies that were used for the experiment are listed in the supplementary table 2. The bands will be analyzed by optical densitometry. The results will be normalized by the expression of β-actin.

### 2.8. Determination of ROS by lucigenin chemiluminescence assay

The chemiluminescent probe lucigenin (bis-N-methylacridinium nitrate) was used to determinate ROS generation in thoracic aorta. In brief, the aortas were frozen in liquid nitrogen and homogenized in phosphate buffer. 50µL of the homogenate with 175 µL of lucigenin (0.005 mmol/L) and assay buffer were transfered to a White 96-well microplate. A luminometer (FlexSation 3 microplate reader (Molecular Devices, San Jose, USA) was used to read the plate. After the first reading, the substrate for the NADPH oxidase enzyme was added (NADPH (0.1 mmol/L) to the suspension and a second read of the plate was obtained. Results are shown as relative light units (RLU)/mg of protein. To confirm that ROS signal is being generated by mitochondria, some pieces of aorta were incubated with a mitochondria-targeted antioxidant agent (mitotempo, 1 x 10^-4^ mmol/L, for 1h).

### 2.9. Determination of H_2_O_2_ by Amplex Red

The aortic concentration of H_2_O_2_ was determined through the fluorogenic substratedAmplex ® Red (#A22188, Invitrogen, Carlsbad, CA, USA). Amplex red reacts with H_2_O_2_ to produce fluoroscentresofurin. The concentration of H_2_O_2_ was measured in a fluorometer (FlexSation 3 microplate reader (Molecular Devices, San Jose, USA). Results were expressed as nmol/L/mg protein.

### 2.10. Determination of mitophagy in fresh thoracic aortic

To analyze whether ethanol impairs mitophagy process in thoracic aorta, we treated a unique mouse model to track mitophagy in fresh tissue, mt-Keima mouse (a gift from Dr. Finkel at University of Pittsburgh). Keima is a coral protein, genetically inserted in the mitochondria, it is a pH-sensitive and resistance to lysosomal proteases. At the physiological pH of the mitochondria (pH 8.0), the shorter-wavelength excitation predominates (green), whereas that within the acidic lysosome (pH 4.5), mt-Keima undergoes a gradual shift to a longer-wavelength excitation (red)^41^. After the treatments described above, aortae were harvested freely of PVAT and longitudinally open. Then, they were placed on the glass slides with endothelium upward and covered with a coverslip. Mitochondria track was analyzed in an Revolve Echo Motorized Fluorescence microscopy and for quantification of fluorescence intensity of the red and green signs was used Image J software. The images were converted to black and white, and the regions of interest was selected 40 times by tracing using a rectangle selection tool in different places per image. The mean was calculated, and the ratio (red/green) was devised to assess the mitophagy index.

### 2.11. Statistical analysis

One-way analysis of variance (one-way ANOVA) and Two-way analysis of variance (two-way ANOVA) followed Tukey post-test was performed to detect differences between the values under study. The concentration-response curves were fitted by nonlinear regression analysis. Analyses were performed using Prism software (GraphPad Prism, San Diego, CA, USA). A difference was considered statistically significant when P ≤ 0.05.

## 3. RESULTS

### Ethanol-induced hypercontractility relies on endothelium integrity and RASS activation

We first found that ethanol reduced food and water intakes, which were not affected by losartan (Suppl. Fig. 1A and B). Furthermore, ethanol treatment impaired the body weight again, while losartan induced a further body weight gain in control and ethanol groups (Suppl. Fig. 1C). To analyze whether the animals received appropriate dose of losartan, we measured food intake and multiplied by the amount of losartan incorporated into food. As expected, mice under losartan treatment received 10.2± 0.5 grams per day/Kg of drug. Since ethanol consume has been associated with cardiac, hepatic, and renal changes, at the end of 12 weeks of treatment we measured the weights of those organs. Ethanol did not change heart weight, but increased the hepatic and renal mass, while losartan protected from hepatic abnormalities associated with ethanol treatment (Suppl. Fig. 1D-F).

The vascular function is orchestrated by three major compartments endothelium, vascular smooth muscle cells (VSMC), and PVAT, which is mostly composed by adipocytes^42,43^. By working with aortic rings with intact (Endo+) or denuded endothelium (Endo-) and with (PVAT+) or without (PVAT-) PVAT, we investigated by which vascular compartment ethanol can lead to vascular dysfunction. Ethanol treatment increased vascular contractility to phenylephrine in Endo+ aortic rings (Fig. 1A and table 1), which was blunted by endothelium removal (Fig. 1B and table 1). Furthermore, the presence of PVAT produced an anti-contractile effect (reduced contractility and alterations in pD_2_ for phenylephrine) in aortic rings with and without endothelium from control and ethanol treatment groups (Fig. 1A and 1B and table 1), suggesting that ethanol is leading to vascular dysfunction in an endothelium dependent manner with no interference of PVAT. We next interrogated whether ethanol treatment affects the aortic contractility to an additional constrictor agent (KCl). We did not find any changes for KCL-induced contraction (Fig. 1C and 1D).

**Figure 1.**
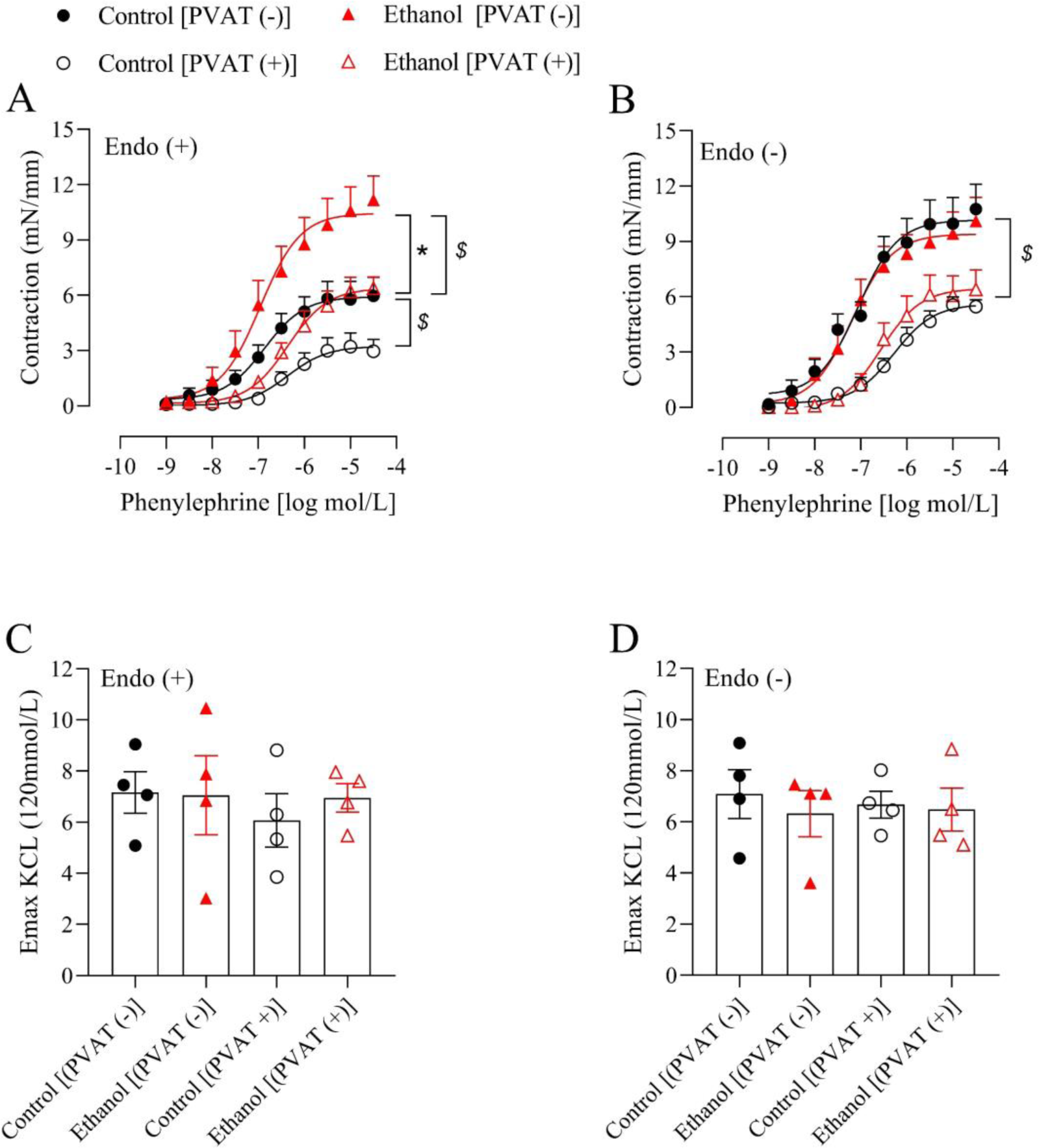
Ethanol consumption increases vascular contractility in an endothelium-dependent manner without PVAT effects. Concentration responses curves to phenylephrine in aortic rings with [Endo (+)] (A) or without endothelium [Endo (-)] (B), in presence [PVAT (+)] or absence of [PVAT (-)] and KCL (120mmol/L)-induced vascular contractility in the same aortic rings (C and D) from mice treated with vehicle or ethanol for 12 weeks. Values are represented as the mean ± SEM of n=4-6 animals per group. *P<0.05 vs control group; ^$^P<0.05 vs PVAT (+). Statistic analyze was performed by a two-way ANOVA followed by the Tukey post-test.

**Table 1:**
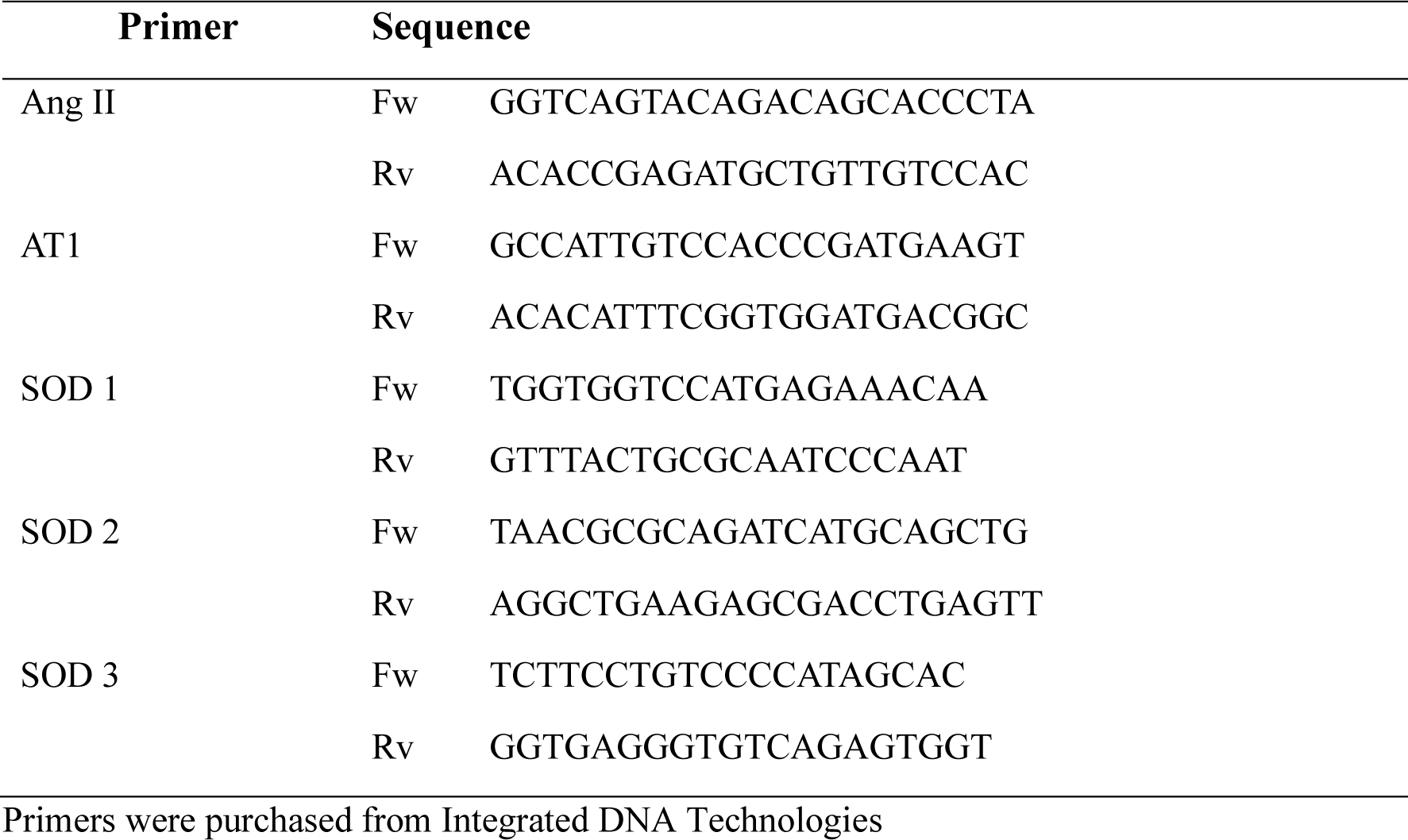
List of primers.

RAAS is placed as an important effector of the cardiovascular injury induced by ethanol^8^. Thus, we evaluated the participation of Ang II in ethanol-induced vascular dysfunctionby treating mice with losartan (10 mg/kg). Blockade of AT1 receptors prevented ethanol-induced hypercontractility to phenylephrine in aortas with intact endothelium (Fig. 2A and table 1) Furthermore, via RT-PCR, we found that chronic ethanol consumption increased mRNA expression of Ang II in thoracic aorta (Fig. 2B) but did not change mRNA expression of AT1 receptor (Fig. 2C), suggesting that RAAS participates of ethanol-induced vascular dysfunction. In addition, we did not observe any effects on vascular remodeling determined by wall thickness (Fig. 2D). Since, no change on PVAT or VSMC function were detected, as well as on vascular structure, for the subsequent experiments, we focused solely on vascular contractility in arteries with intact endothelium.

**Figure 2.**
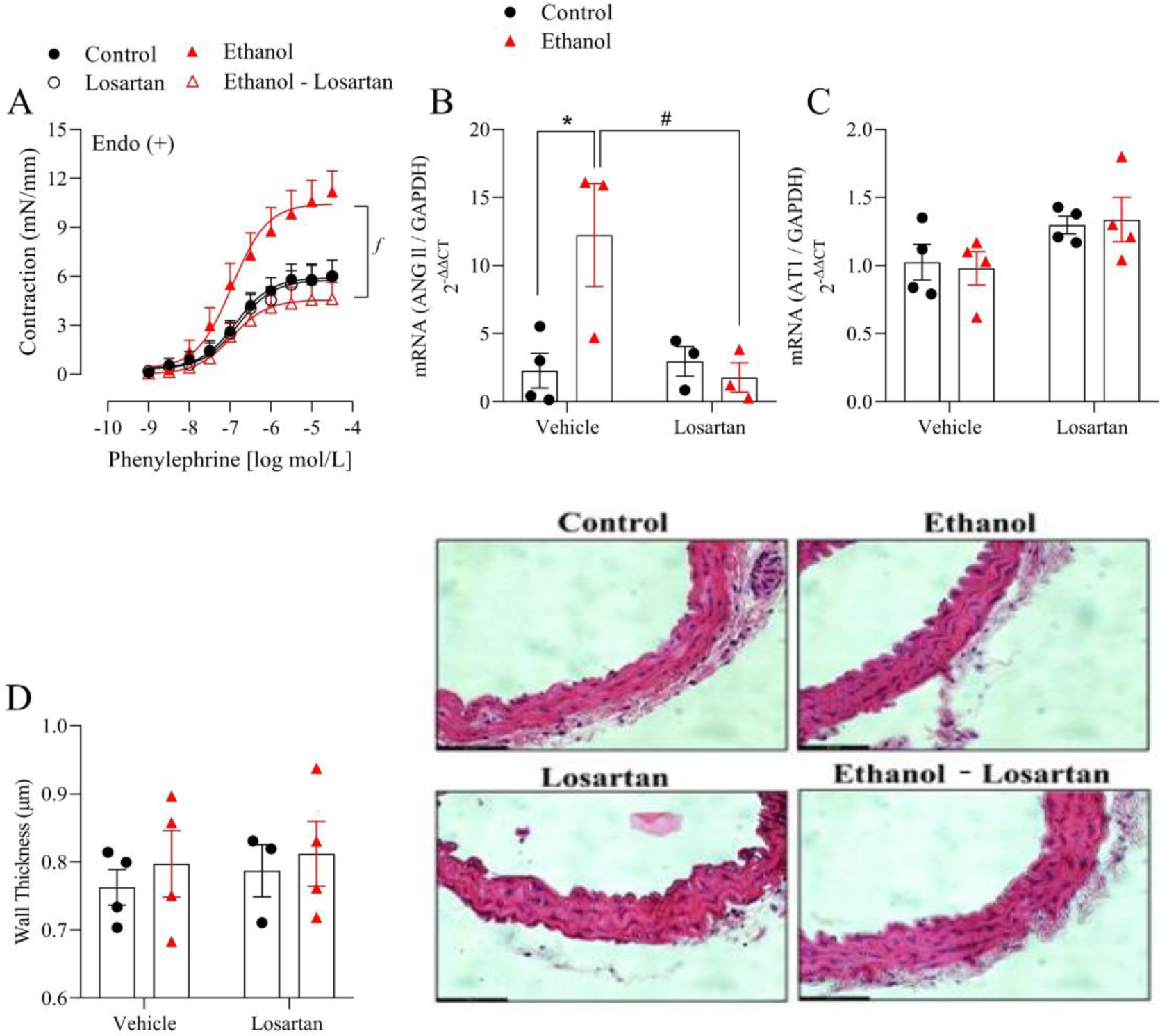
Ethanol-induced hypercontractility depends on Ang II signaling. Concentration responses curves to phenylephrine in thoracic aorta with intact endothelium [Endo (+)] (A). Angiotensin II and AT1 gene expression (B and C) determined by RT-PCR. Wall thickness measured in aortas stained with hematoxylin & eosin (H&E) (D) from mice maintained for 12 weeks on normal or losartan (10 mg/kg) diets and treated with vehicle or ethanol for 12 weeks. Ruler length: 62.3um.Values are represented as the mean ± SEM of n=3-8 animals per group. ^ƒ^P<0.05 vs ethanol-losartan group; *P<0.05 vs control group; ^#^P<0.05 vs ethanol-losartan group. Statistic analyze was performed by a two-way ANOVA followed by the Tukey post-test.

**Table 1a.** Emax (mN) and pD2 values of phenylephrine-induced contraction in thoracic aorta of control, ethanol, Losartan and ethanol - Losartan (Eth-Los) groups.

| Groups | Phenylephrine [PVAT (+)] |  | Phenylephrine [PVAT (-)] |  |
| --- | --- | --- | --- | --- |
|  | E<sub>max</sub> (mN) | pD<sub>2</sub> | E<sub>max</sub> (mN) | pD<sub>2</sub> |
| Control (Endo +) | 2.9 ± 1.4 (5) | 6.4 ± 0.2 | 5.9 ± 2.7 (7)<sup>a</sup> | 6.9 ± 0.2 |
| Ethanol (Endo +) | 6.4 ± 1.7 (8)<sup>ac</sup> | 6.4 ± 0.1 | 11.2 ± 3.2 (6)<sup>ab</sup> | 6.9 ± 0.2 |
| Control (Endo -) | 5.5 ± 0.4 (7) | 6.3 ± 0.1 | 10.8 ± 1.4 (5)<sup>a</sup> | 7.1 ± 0.2<sup>a</sup> |
| Ethanol (Endo -) | 6.4 ± 1.1 (8)<sup>bc</sup> | 6.6 ± 0.2<sup>c</sup> | 10.1 ± 1.3 (8)<sup>a</sup> | 7.2 ± 0.2<sup>a</sup> |
|  | Phenylephrine [ PVAT (+)] |  | Phenylephrine [ PVAT (-)] |  |
|  | E<sub>max</sub> (mN) | pD<sub>2</sub> | E<sub>max</sub> (mN) | pD<sub>2</sub> |
| Control (Endo +) | 2.9 ± 1.4 (5) | 6.4 ± 0.2 | 5.9 ± 2.7 (7) | 6.9 ± 0.2 |
| Losartan (Endo +) | 2.9 ± 0.5 (8) | 6.4 ± 0.2 | 6.0 ± 0.9 (8) | 6.8 ± 0.2 |
| Ethanol (Endo +) | 6.4 ± 1.7 (8)<sup>d</sup> | 6.4 ± 0.1 | 11.2 ± 3.2 (6)<sup>d</sup> | 6.9 ± 0.2 |
| Eth+Los (Endo +) | 2.4 ± 0.7 (8) | 6.2 ± 0.3 | 4.6 ± 1.0 (6) | 6.9 ± 0.2 |

Values represent the mean ± SEM of the different experimental groups; the numbers in parentheses represent the “n” of each experimental group; <sup>a</sup>Difference in relation to the control [PVAT (+)]; <sup>b</sup>Difference in relation to control [PVAT (-)]; <sup>c</sup>Difference in relation to ethanol [PVAT (-)]; <sup>d</sup>Difference in relation to the other groups (p<0.05; two-way ANOVA followed by the Bonferroni post-test).

### Ethanol impairs vascular mitochondria biogenesis, dynamic, and mitophagy dependent on RAAS activation

To determine the mechanism by which Ang II promotes vascular dysfunction in mice under chronic ethanol treatment, we firstly evaluated aortic mitochondrial biogenesis and dynamic. We found that PGC1-α, a marker of biogenesis, and MTF2, an essential protein for fusion, were suppressed in aortas from ethanol-treated mice (Fig. 3A and 3B). Although, we did not see a statistical difference in optic atrophy type 1 (OPA-1), a marker of mitochondria (Fig. 3C). Losartan treatment protected from ethanol-induced suppressed mitochondria fusion.

**Figure 3.**
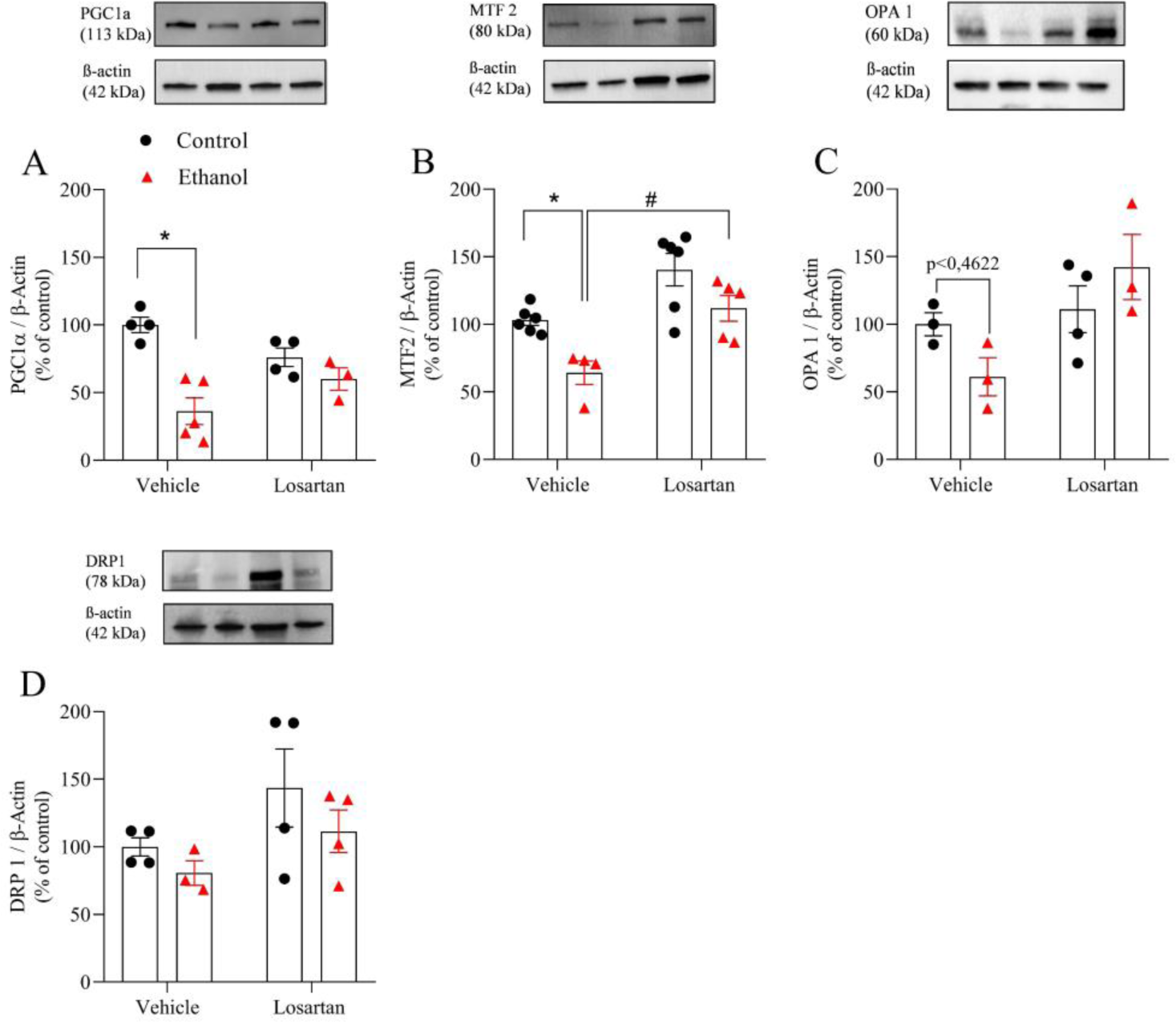
Ethanol decreases fusion (MTF2) dependent on Ang II signaling and mitochondrial biogenesis. Protein expression of PGC1α (A), MTF2 (B), OPA1 (C) and DRP1(D) determined by immunoblotting in thoracic aorta of mice maintained for 12 weeks on normal or losartan (10 mg/kg) diets and treated with vehicle or ethanol for 12 weeks. Values are represented as the mean ± SEM of n=3-6 animals per group. *P<0.05 vs control group; ^#^P<0.05 vs ethanol - losartan group. Statistic analyze was performed by a two-way ANOVA followed by the Tukey post-test.

Next, we examined whether mitophagy, a process to recycle defective mitochondria and important on regulating the vascular physiology^44^, is disturbed by ethanol treatment. We observed that ethanol treatment significantly suppressed aortic Pink-1 expression and LC3A/B ratio (Fig. 4A and 4C), but not Parkin (Fig. 4B), although a strong trend was observed. Interestingly losartan prevented ethanol-induced disruption of mitophagic flux (Fig. 4A and 4C). By utilizing an elegant mouse model to track mitophagy in fresh tissues, the mt-Keima mouse ^41^, we confirmed that ethanol impaired mitophagy flux via RASS overactivation (Fig. 4D) by finding a reduced mitochondria content in the lysosome compartment, which was blunted by AT1 receptor antagonist treatment. Overall, these data suggest that ethanol leads to a dysfunctional mitochondrion likely by disturbing mitochondria dynamic, and recycling via Ang II signaling.

**Figure 4.**
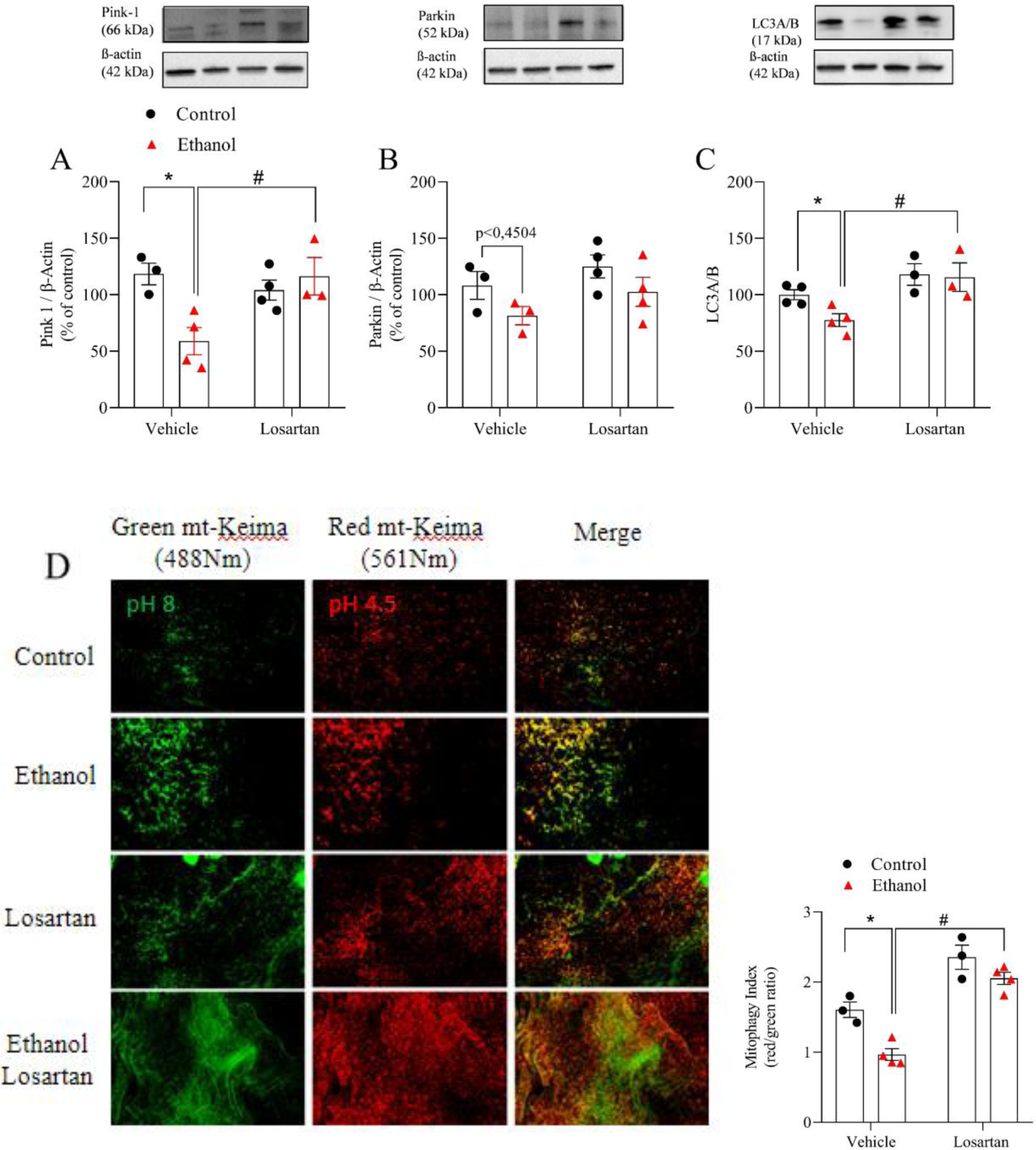
Ethanol decreases mitophagy dependent on Ang II signaling. Protein expression of Pink1 (A), Parkin (B) and LC3AB (C) determined by western immunoblotting and mtKeima red (561 nm excitation) and green (488 nm excitation) signal (D) assessed by fluorescence microscopy in thoracic aorta and the ratio 561:488 (mitophagy index). Data were collected from mice maintained for 12 weeks on normal or losartan (10 mg/kg) diets and treated with vehicle or ethanol for 12 weeks. Values are represented as the mean ± SEM of n=3-4 animals per group. *P<0.05 vs control group; ^#^P<0.05 vs ethanol - losartan group. Statistic analyze was performed by a two-way ANOVA followed by the Tukey post-test.

### mtROS is involved in ethanol-induced hypercontractility dependent on Ang II signaling

Generally, mitochondria dysfunction is associated with exacerbated mtROS generation^15^. Thus, we investigated whether ethanol leads to increase in mtROS in aorta via Ang II signaling. We observed that ethanol treatment increased ROS formation, measured by lucigenin chemiluminescence assay, which was abolished by losartan treatment. To confirm that this signal is mtROS, some aortae were incubated with mitotempo (1 x 10^-4^ mmol/L, mitochondria-targeted antioxidant agent) prior lucigenin assay. Mitotempo treatment suppressed ROS formation in ethanol group and did not affect the other groups, suggesting that RAAS activation is leading to ethanol-induced mtROS generation (Fig. 5A). Via RT-PCR, we found that chronic ethanol consumption did not change expression of SOD1, SOD2 and SOD3 (Fig. 5C-E), but it did decrease the SOD2 protein expression, whereas blockade of AT1 receptors prevented such change (Fig. 5B). Next, we examined whether ethanol triggers vascular dysfunction via mtROS by incubating thoracic aortas with MnTMPyP (0.03 mmol/L, mitochondrial antioxidant agent) and carbonyl cyanide m-chlorophenyl hydrazine (CCCP, 10^-3^ mmol/L, mitochondrial uncoupler). Both agents abrogated the hypercontractility to phenylephrine induced by ethanol treatment (Fig. 5F and 5G and table 2) and did not change the vascular response in the groups treated with losartan (Fig 5H and 5I and table 2).

**Figure 5.**
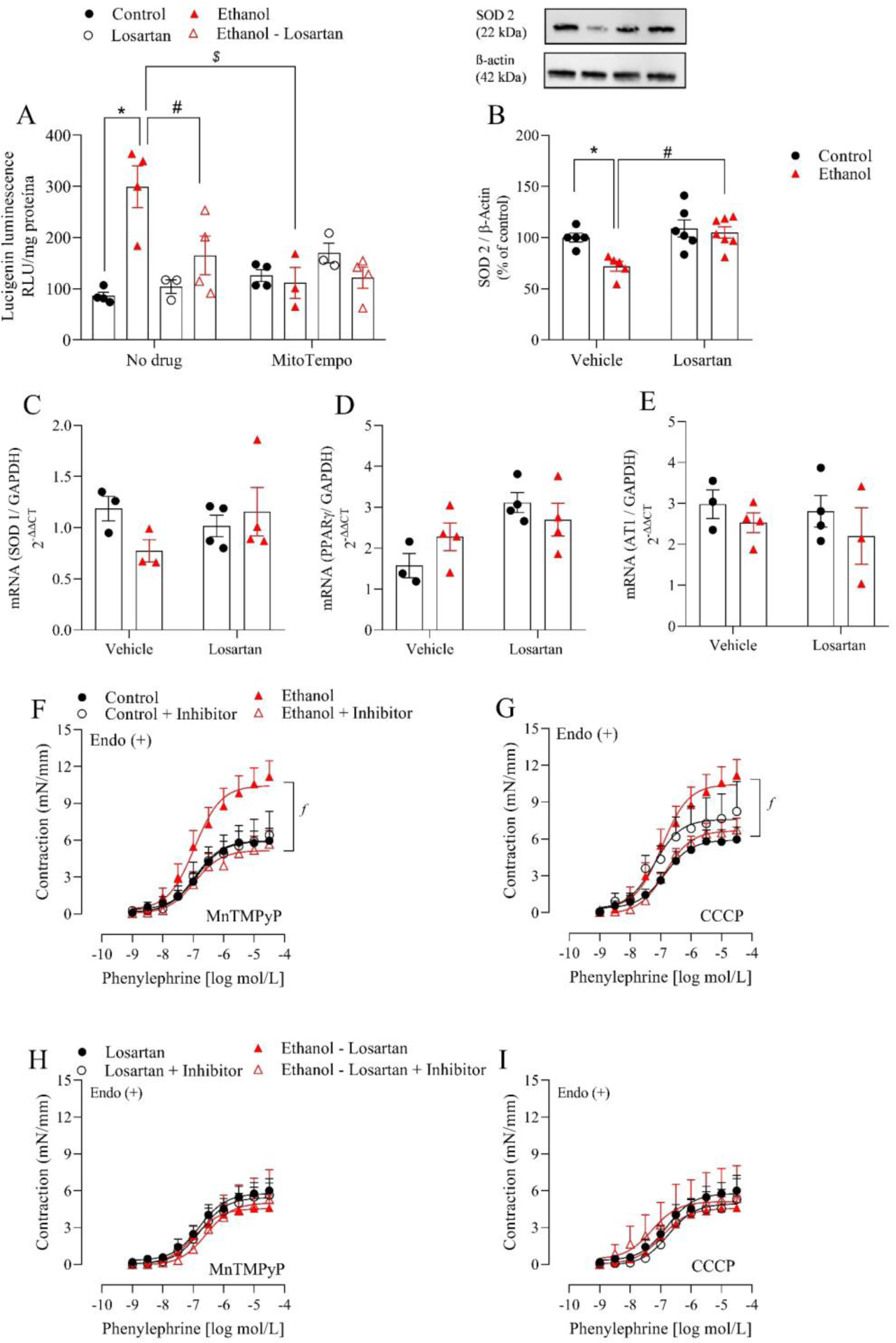
Mitochondria derived ROS drives ethanol-induced hypercontractility dependent on Ang II signaling. ROS generation determined by lucigenin chemiluminescence assay in thoracic aorta with or without mitotempol (1 x 10^-4^ mmol/L, mitochondria-targeted antioxidant agent) (A). Protein expression of SOD2 (B) determined by western immunoblotting, and SOD1 (C), SOD2 (D), and SOD3 (E) gene expression determined by RT-PCR. Concentration responses curves to phenylephrine in thoracic aorta with [Endo (+)]in presence or absence of MnTMPyP (0.03 mmol/L) (F and H) or CCCP (10^-3^ mmol/L) (G and I). Data were collected in thoracic aorta of mice maintained for 12 weeks on normal or losartan (10 mg/kg) diets and treated with vehicle or ethanol for 12 weeks. Values are represented as the mean ± SEM of n=3-8 animals per group. *P<0.05 vs control group; ^#^P<0.05 vs ethanol - losartan group; ^ƒ^P<0.05 vs aorta treated with MitoTempol or MnTMPyP or CCCP. Statistic analyze was performed by a two-way ANOVA followed by the Tukey post-test.

**Table 2:** List of antibodies.

| Antibody | Catalog number | Company | Concentration |
| --- | --- | --- | --- |
| PGC-1α | ab176328 | Abcam | 1:1000 |
| Mitofusion-2 (MTF2) | 9482 | Cell siganling | 1:1000 |
| OPA-1 | 80471 | Cell siganling | 1:1000 |
| DRP1 | 5391 | Cell siganling | 1:1000 |
| PINK-1 | ab23707 | Abcam | 1:1000 |
| Parkin (Prk8) | ab77924 | Abcam | 1:1000 |
| LC3A/B | 4108 | Cell siganling | 1:1000 |
| SOD2 | 13141 | Cell siganling | 1:1000 |
| Catalase | 12980 | Cell siganling | 1:1000 |
| B-actin | A3854 | Sigma | 1:20000 |

### Decreases in aortic H_2_O_2_ and NO via Ang II signaling contributes to ethanol-induced vascular dysfunction

Reduced SOD2 expression can increase O_2_^•-^, which in turn affects H_2_O_2_ levels and inactive the vasodilatory effects of NO^45,46^. Here we observed that chronic ethanol consumption decreased aortic H_2_O_2_ levels, while losartan treatment prevented this alteration (Fig.6A). No changes in catalase protein expression were observed (Fig. 6B). To examine whether decreased H_2_O_2_ is affecting the vascular contractility, thoracic aortas were incubated with EUK-134 (0.01 mmol/L), synthetic superoxide dismutase and catalase mimetic) prior phenylephrine curves. We observed that EUK-134 increased vascular contraction in control group, but not in ethanol group (Fig. 6C and table 2). EUK-134 also increased vascular contraction in losartan-treated mice, suggesting that losartan prevented the loss of H_2_O_2_, which might be acting as a vasodilatory molecule (Fig. 6D and table 2). We next determined whether suppressed NO signaling might be associated with ethanol-induced vascular dysfunction. With this purpose, we inhibited nitric oxide synthase (NOS) with L-NAME (0.1 mmol/L), which demonstrated that NOS inhibition increased the phenylephrine response in control, losartan, and ethanol-losartan groups, but not in ethanol group (Fig. 6E and table 2), suggesting that ethanol reduces NO bioavailability via RASS activation (Fig. 6F and table 2).

**Figure 6.**
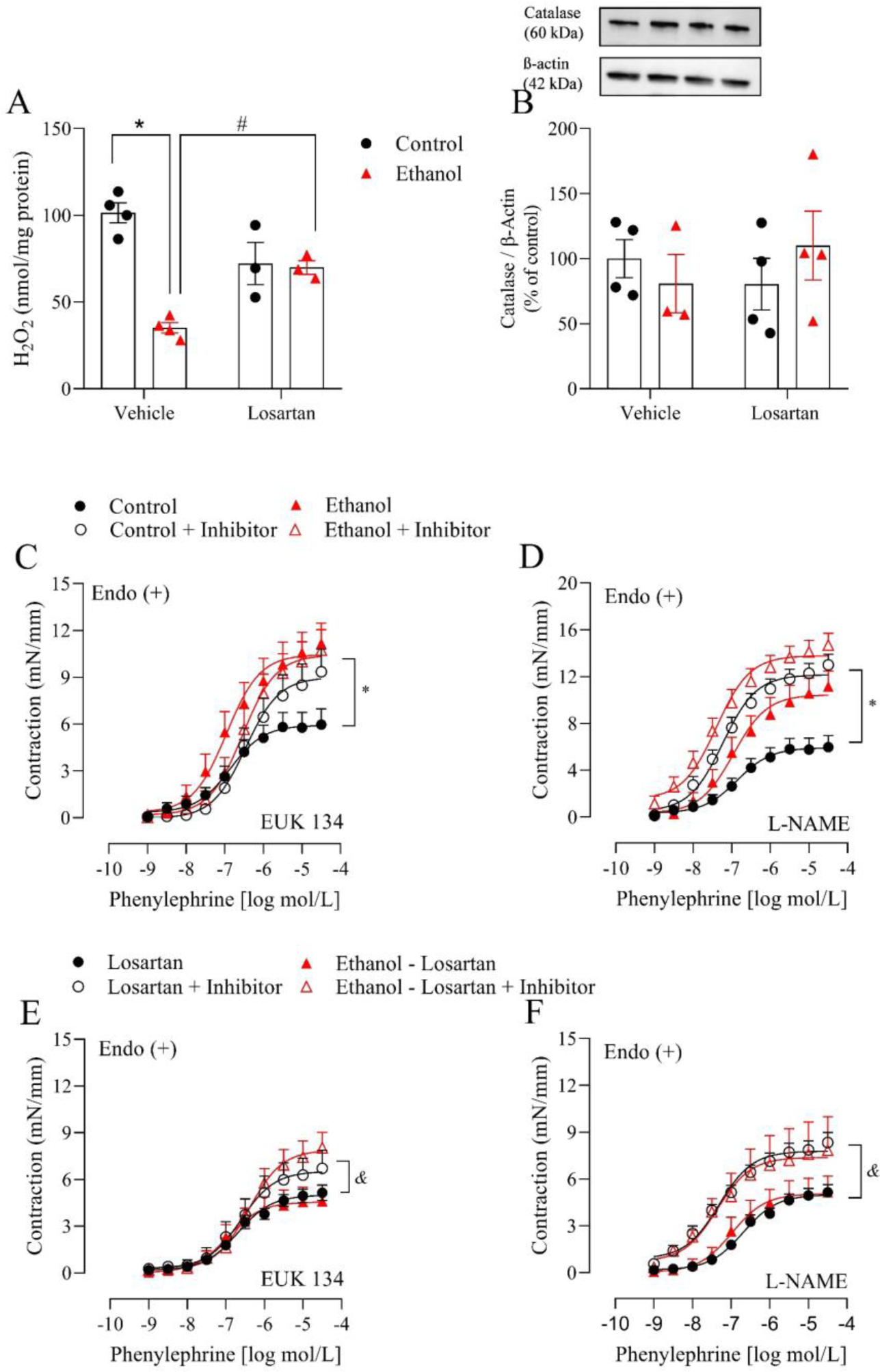
Ethanol induces vascular dysfunction by decreasing H_2_O_2_ and nitric oxide bioavailability dependent on Ang II signaling. Aortic H_2_O_2_ (A) determined by Amplex Red and aortic protein expression of catalase (B). Concentration responses curves to phenylephrine in thoracic aorta with [Endo (+)] in presence or absence of EUK 134 (0.01 mmol/L) (C and D) or L-NAME (0.1 mmol/L) (E and F). Data were collected in thoracic aorta of mice maintained for 12 weeks on normal or losartan (10 mg/kg) diets and treated with vehicle or ethanol for 12 weeks. Values are represented as the mean ± SEM of n=3-8 animals per group. *P<0.05 vs control group; ^#^P<0.05 vs ethanol - losartan group; ^&^P<0.05 vs presence of EUK134 or L-NAME. Statistic analyze was performed by a two-way ANOVA followed by the Tukey post-test.

**Table 2a.**
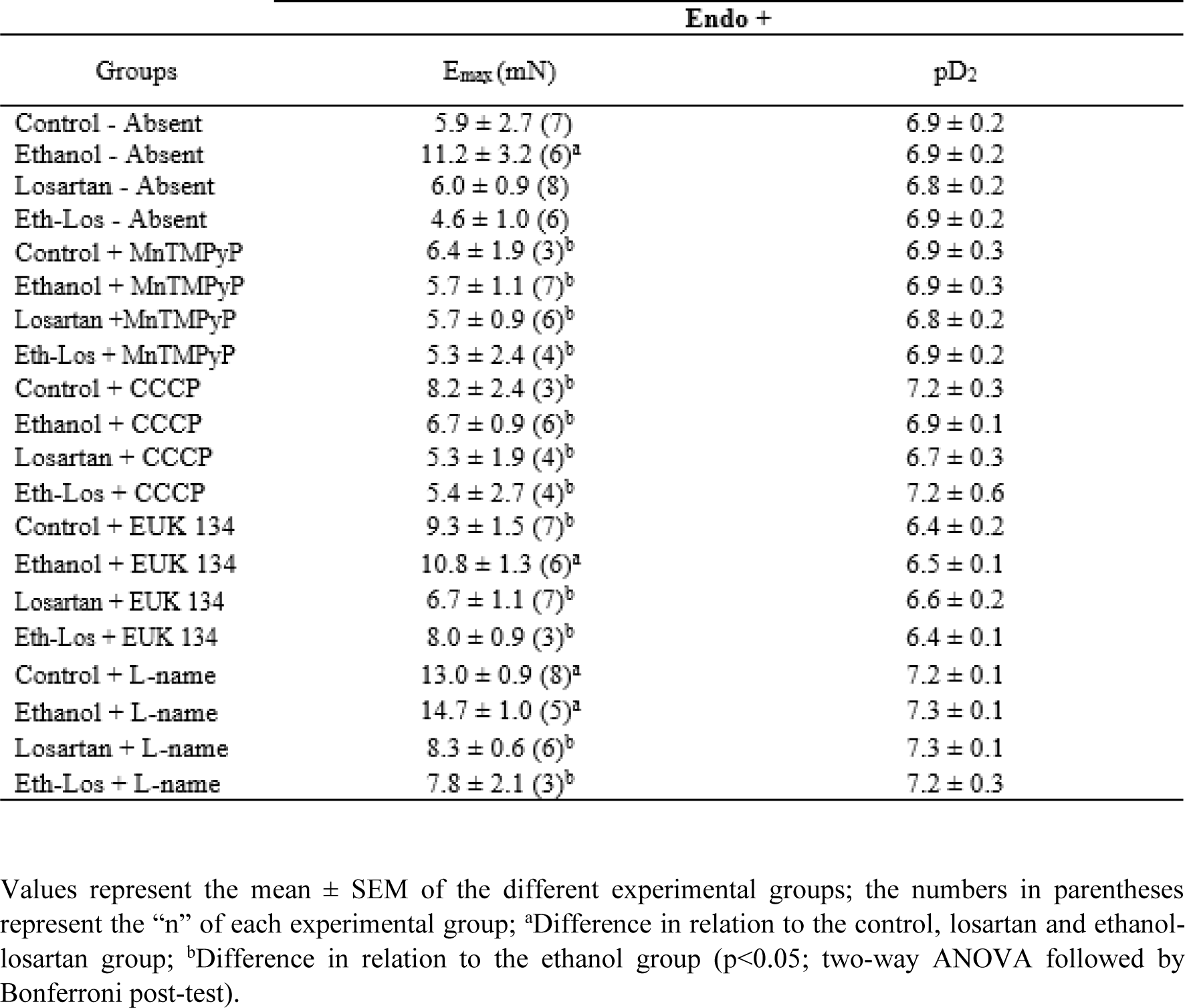
Emax (mN) and pD2 values of phenylephrine-induced contraction in thoracic aorta of control, ethanol, losartan and ethanol-losartan (Eth-Los) groups with or without inhibitor.

## 4. Discussion

Although the mechanism of vascular dysfunction induced by high ethanol consumption has been extensively studied, little is known about the importance of mitochondria on regulating this cardiovascular risk. Mitochondria dysfunction has been described as a crucial mechanism in the genesis of multiple diseases including cardiovascular^47,48^. Therefore, elucidating if disturbs in mitochondria quality is a trigger for ethanol-induced vascular injury is paramount interest. In this novel study, we are confirming our hypothesis that ethanol induces vascular dysfunction by impairing mitochondria function via RAAS activation. As main results we are describing that ethanol consumption leads to vascular dysfunction by impairing mitochondria biogenesis, dynamic, and recycle. As a result of mitochondrial dysfunction, ethanol treatment augmented mtROS formation and suppressed aortic NO and H_2_O_2_ levels, such effects seemed to be dependent on RAAS activation, since AT1 receptors blockade prevented vascular deleterious effects of ethanol.

Chronic ethanol consumption affects cardiac, hepatic, and urinary systems, as well as food and water intake in humans and rodents^49–54^. Therefore, we measured food and water intake, as well as liver, kidney, and heart masses. Here, we observed a reduction in food and liquid intake in animals treated with ethanol and a consequent reduction in body weight gain followed by increase in liver and kidney weights, but no alteration in heart among the groups. In humans ethanol impairs intestinal transport and absorption of nutrients ^55^, while the liver may suffer hypertrophy due to the accumulation of water, proteins, and fatty acids ^56^ and kidneys undergo dysregulated hydric balance compromising theirs structure^57^. Thus, our data suggest that the protocol used in our study represents an appropriate method to study alcoholism and cardiovascular risk, since it mimics multiple features found in humans.

Changes in vascular contractility have been reported before in models of ethanol treatment^4,5,58–62^, however the molecular and cellular mechanisms are not fully elucidated. Herein, we observed that ethanol treatment does not induce vascular remodeling, but it does increase vascular contractility in an endothelium dependent manner. We further demonstrated that PVAT is not dysfunctional after 12 weeks of ethanol treatment. Previous findings from our group demonstrated that ethanol treatment for 12 weeks affects the anti-contractile effects of PVAT of mesenteric arteries^4^, but not aortic PVAT^35^. Whereas PVAT in mesenteric arteries is mostly formed by white adipocytes, PVAT from thoracic aorta is predominantly composed by brown adipocytes, which can interfere on the communication with its neighboring vasculature by producing a different spectrum of adipokines, anti-contractile molecules, and extracellular vesicles^63^. Furthermore, in acute ethanol treatment PVAT exerts protective effect against vascular dysfunction ^64^. Thus, duration of ethanol treatment and type of PVAT studied must be considered in research focusing on the effects of ethanol on vascular function.

A relevant participation of RAAS in ethanol-induced cardiovascular alterations has been demonstrated before^5,7,8,22,23^. For this reason, RASS is placed as an important effector of the cardiovascular actions induced by ethanol. In our study, we confirmed the participation of RAAS in vascular dysfunction in ethanol treated mice by demonstrating that losartan prevented the increase in vascular contractility induced in mice chronically treated with ethanol. In addition, we observed an increase in the expression of the Ang II gene in the thoracic aorta from ethanol group, but with no changes in the expression of AT1 receptors. In the classic RAAS, Ang II is a biologically active peptide produced systemically and locally in the vascular wall^65^. Although, it is well-known that RAAS plays a major role on chronic ethanol consumption-induced vascular injury, the downstream signaling pathways are still not fully explained.

Previous studies have demonstrated that Ang II negatively changes mitochondria homeostasis in the cardiovascular pathophysiology^27,66–68^, but their contribution to the pathophysiology of vascular dysfunction in ethanol treatment is unknown. In an attempt to maintain homeostasis during pathophysiological situation, mitochondria change its morphology, biogenesis, mitophagy, and ROS production ^69,70^. Here, we showed that ethanol impairs mitochondria biogenesis - analyzed by the expression of PGC-1α. PGC-1α is one of the main transcription factors responsible for stimulating mitochondrial biogenesis, a process that involves the formation of new mitochondria by adding proteins and membranes to pre-existing mitochondria^71^. Some studies have shown that ethanol consumption reduces the phosphorylation of cAMP-response element binding protein (p-CREB) in cardiac tissue ^72^ and neuronal cells ^73^. CREB is the main transcription factor that regulates the transcription of the PGC-1α gene^74^. Although we did not measure CREB expression, we can hypothesize that ethanol treatment attenuated the expression or activity of CREB in the vasculature.

Mitochondria are highly dynamic organelles undergoing coordinated cycles of fission and fusion, so called mitochondrial dynamic, which is fundamental for the organelle function ^75^. We found that MTF2, mitochondrial protein responsible for fusion process, expression was reduced by ethanol consumption with no difference in DRP1 (mitochondria fission inducer), interestingly losartan prevented these changes, suggesting ethanol decreases mitochondria fusion via RASS activation. Studies have shown that Ang II can dysregulate the balance between mitochondrial fusion and fission. For instance, Ang II reduces mitochondria fusion via suppressing MTF2 in cardiomyocytes ^76–79^, which are consistent with our findings.

To maintain a tunned mitochondria function and a proper cellular function, the removal of damaged mitochondria through autophagy (mitophagy) must occur. Disturb in mitophagic flux has been reported in different conditions including aging, neurodegenerative, and cardiovascular diseases^80–83^. In brain, chronic ethanol exposure in adult mice causes impairment of the autophagy-lysosome pathway ^84^, while in esophageal keratinocytes, autophagy mitigates ethanol-induced mitochondrial dysfunction^85^. We found that ethanol decreased mitophagy related proteins (LC3A/b ratio and Pink1), we further observed that mitochondria content in acid pH (lysosome environment) is attenuated in aorta from mt-Keima mice, such changes were dependent on RAAS activation since losartan treatment prevented mitophagy disruption. An interface between suppressed MTF2 and loss of Pink1 and Parkin has been reported before in cardiomyocytes treated with Ang II, which led to accumulation of damaged mitochondria^79^. Therefore, we can suggest that ethanol incudes mitochondria dysfunction in the vasculature via suppressed biogenesis and fusionand impairing mitochondria recycle.

Ang II exerts its potent oxidant effects via AT1 receptor and activation of NADPH oxidase and mitochondria^8,21,27,86–88^. Usually increased in fission, reduced in fusion, impaired oxidative phosphorylation, and defective mitophagy leads to accumulation and overproduction of mtROS^89–92^. In our study, we can postulate that dysfunctional mitochondria led to increased mtROS production, which in turn contributed to the vascular hypercontractility to phenylephrine in ethanol treated mice, since a mimetic of Mn-SOD or a mitochondrial uncoupler prevented ethanol treatment-induced vascular dysfunction. Besides the dysfunctional mitochondria, ethanol decreased SOD2 enzyme protein expression. SOD2 is a major component in the mitochondria that manages ROS levels in the mitochondrial matrix^93,94^. Thus, decrease in SOD2 could be an additional mechanism by which ethanol increased superoxide anion in the vasculature. Interestingly blockage of Ang II signaling protected against ethanol-induced mitochondria dysfunction, decrease in SOD2, and oxidative stress.

Furthermore, ethanol decreased aortic H_2_O_2,_ which also could have occurred by the suppression of SOD2 expression, since SOD2 determines how much superoxide anion is converted into H_2_O_2_. The effects of H_2_O_2_ in the vasculature is concentration dependent ^95,96^, while studies place H_2_O_2_ as a deleterious ROS^90,97,98^, we ^64,99^ and others^100–102^ have found that H_2_O_2_ display vasodilatory effects. For instance, we have demonstrated that acute dose of ethanol increases H_2_O_2_ in PVAT as a protective mechanism for the vasculature^64^. Moreover, the use of EUK-134, a synthetic SOD and catalase mimetic, did not change the vascular response in ethanol treated mice, but it induced hypercontractility in the other groups, suggesting that vascular H_2_O_2_ plays an anti-contractile mechanism, while ethanol treated mice do not respond the same way to EUK-134 because they have suppressed H_2_O_2_ levels. Additionally, we found no difference for catalase (a ubiquitous antioxidant enzyme that degrades hydrogen H_2_O_2_ to water and oxygen). Thus, no difference in catalase expression rules out the idea that reduced H_2_O_2_ depends on increased catalase expression.

Finally, we interrogated whether ethanol triggers vascular dysfunction via NO bioavailability. As a limitation of this study, we did not measure the NO levels, but we inhibited NOS enzymes prior phenylephrine curves, which is a well-established pharmacological approach to study NO signaling. L-NAME incubation increased the maximal response in control, losartan, and ethanol-losartan groups, but did not affect the vascular response in ethanol group suggesting that NO bioavailability or signaling was suppressed by RAAS activation during ethanol consume. It is well-known that Ang II decreases NO bioavailability by inhibiting endothelial NOS activation or reducing NO bioactivity via peroxynitrite formation ^103,104^.

In conclusion, this work offers the first evidence that the molecular mechanisms of ethanol-induced vascular dysfunction are mediated by Ang II signaling and depend on impaired mitochondria biogenesis, dynamic, and recycle, which in turn increases mtROS and decreases NO and H_2_O_2_ bioavailability. Thus, the use of inhibitors of RAAS is a therapeutic approach to regulate mitochondrial function and may confer cardiovascular protection for individuals with disorder associated with abusive ethanol drinking.

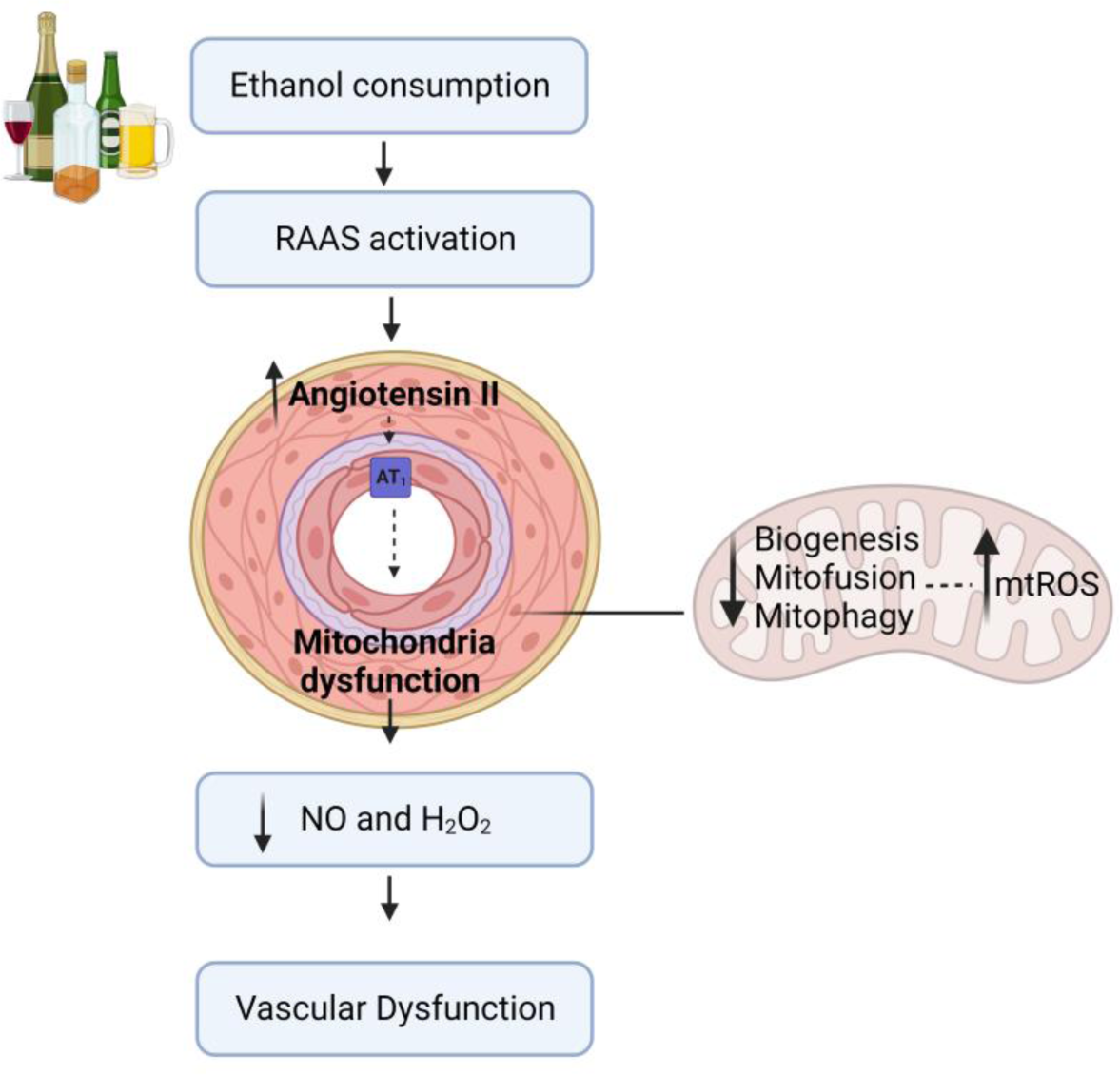

**Supplementary Figure 1.**
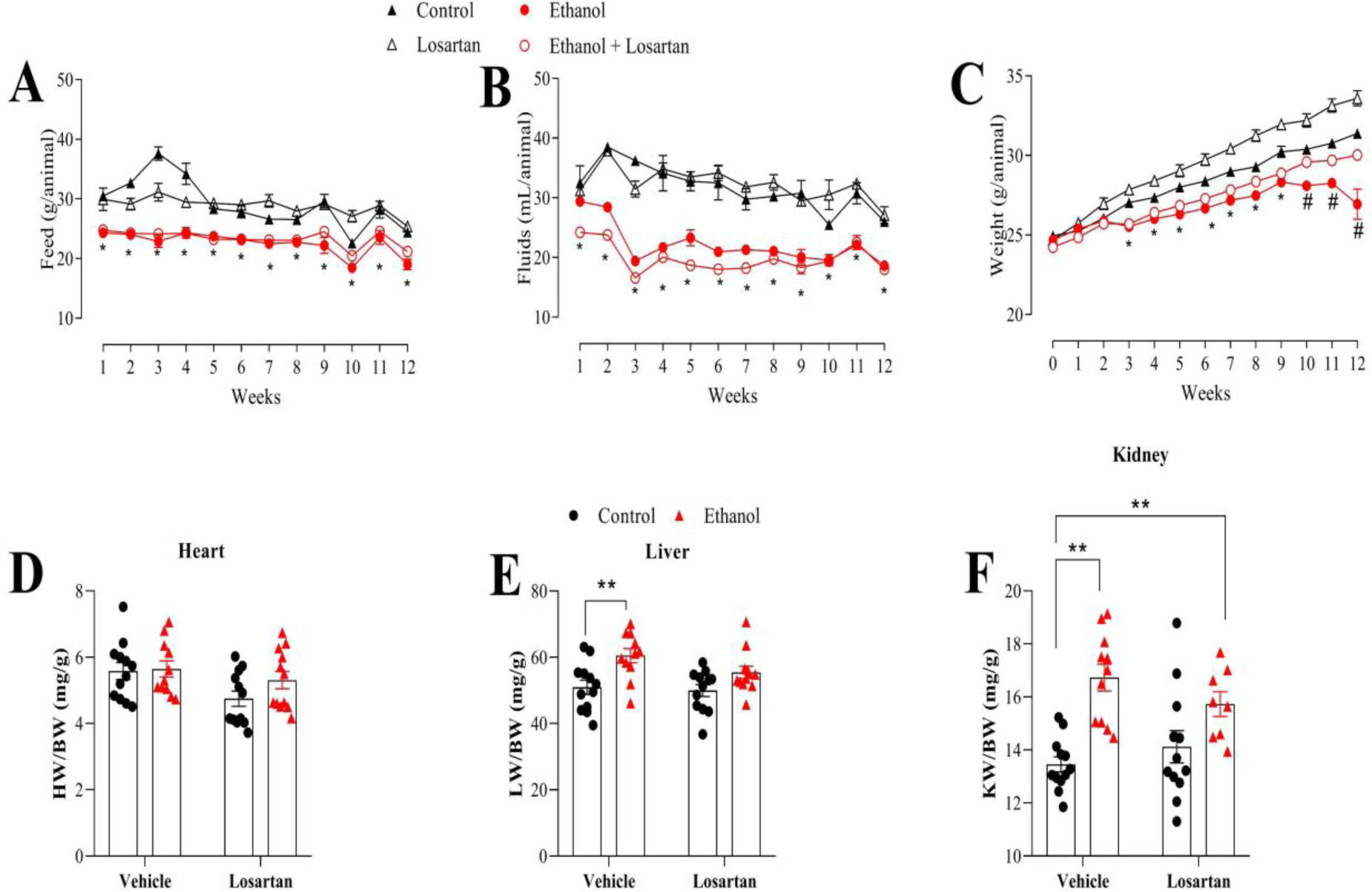
Chronic ethanol consumption promotes change in the nutritional status, weight gain of animals and increases kidney and liver weight in mice. The feed consumption (g) (A), fluid intake (mL)(B), body weight (g) (C), heart weight (mg), liver weight (mg) and kidney weight of mice maintained for 12 weeks on a normal diet or losartan (10 mg/kg), treated with vehicle or ethanol. Values are represented as the mean ± SEM of n=8-16 animals per group. *Difference in relation to the control and losartan groups; *Difference in relation to the control group; (p<0.05; two-way ANOVA followed by the Tukey post-test).

**Supplementary Figure 2.**
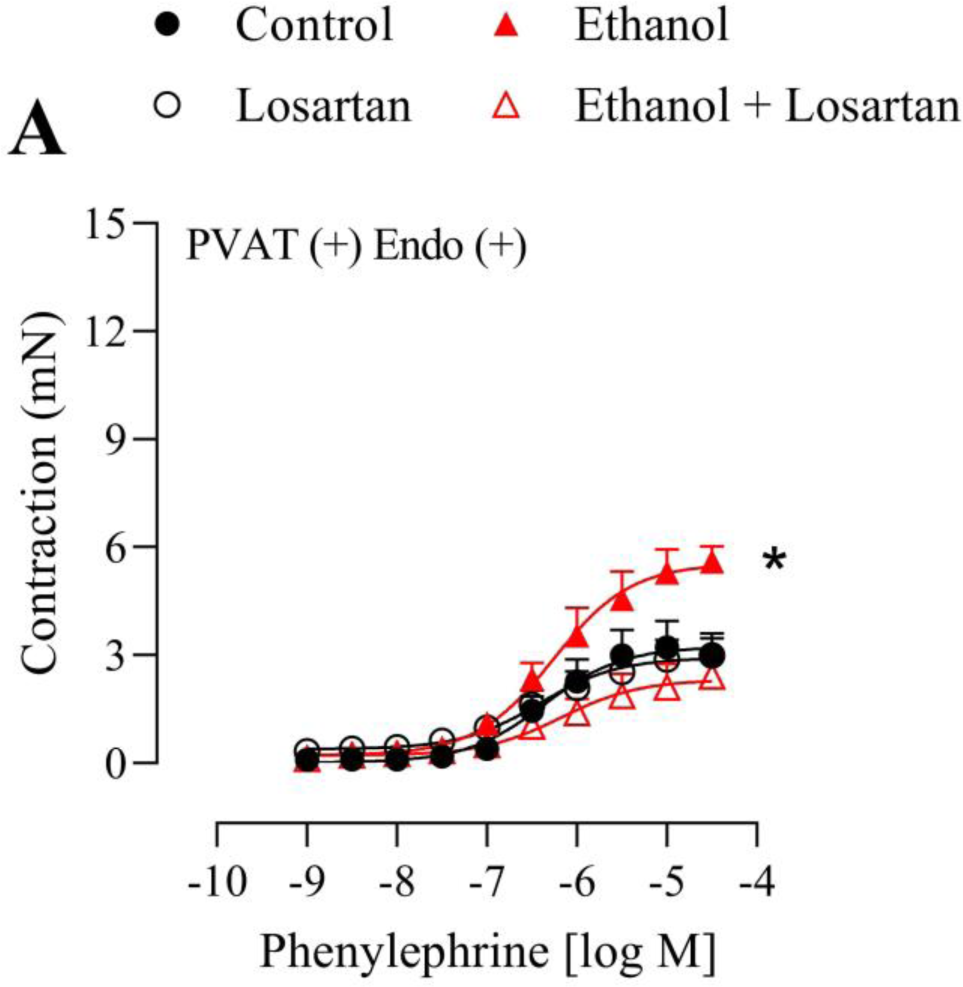
Ethanol-induced hypercontractility depends on Ang II signaling. Concentration responses curves to phenylephrine in thoracic aorta in the presence of endothelium [Endo (+)] and PVAT (A) of mice maintained for 12 weeks on a normal diet or losartan (10 mg/kg), treated with vehicle or ethanol. Values are represented as the mean ± SEM of n=5-8 animals per group. *Difference in relation to the other groups (p<0.05; two-way ANOVA followed by the Tukey post-test).

**Supplementary Figure 3.**
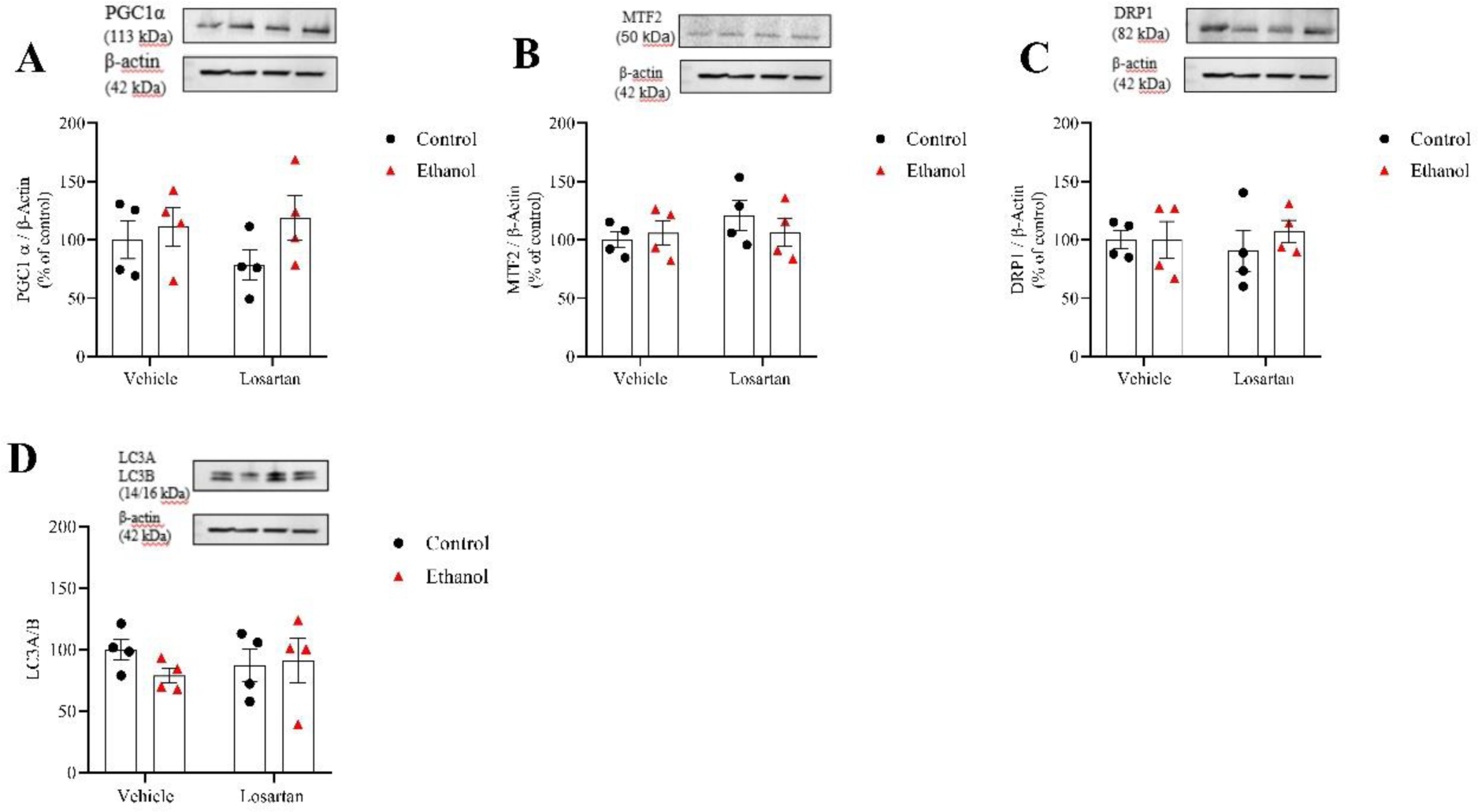
Ethanol does not alter mitochondrial biogenesis, fusion, fission and mitophagy in PVAT. Expression of PGC1α (A), MTF2 (B), DRP1 (C) and LC3A/B (D) determined by western immunoblotting in PVAT of mice maintained for 12 weeks on a normal diet or losartan (10 mg/kg), treated with vehicle or ethanol. Values are represented as the mean ± SEM of n=4 animals per group. (p<0.05; two-way ANOVA followed by the Tukey post-test).

## Acknowledge

Graphical abstract figure was designed in BioRender under license AJ25R0AFAX.

## Funding

Fundação de Amparo à Pesquisa do Estado de São Paulo, Brazil (FAPESP, #2021/13075-5) to WMCA. NHLBI-R00 (R00HL14013903), AHA-CDA (CDA857268), Vascular Medicine Institute, the Hemophilia Center of Western Pennsylvania Vitalant, in part by Children’s Hospital of Pittsburgh of the UPMC Health System, and startup funds from University of Pittsburgh to TBN.

## Disclosure

The authors declare that they have no known competing financial interests or personal relationships that could have appeared to influence the work reported in this paper.

### Abbreviations

NOX: [NADPH] oxidase
ATP: Adenosine triphosphate
Ang II: Angiontensin II
Anti-Drp1: Anti-Dynamin-related protein 1
anti-Mfn2: Anti-mitofusin 2
anti-PGC1α: Anti-Peroxisome proliferator-activated receptor-gamma coactivator (PGC)-1alpha
anti-Pink1: Anti-PTEN Induced Kinase
BAT: Brown adipose tissue
p-CREB: Camp-response element binding protein
CCCP: Carbonyl cyanide m-chlorophenyl hydrazine
cDNA: Complementary DNA
[Endo (+)]: Endothelium
[Endo (-)]: Endothelium denuded
NADPH: Enzyme β-nicotinamide adenine dinucleotide phosphate
EUK-134: Ethylbisiminomethylguaiacol manganese chloride
H&E: Hematoxylin and eosin
MnTMPyP: Manganese (III) tetrakis(1-methyl-4-pyridyl)porphyrin
eMAX: Maximal effect
mtROS: Mitochondrial reactive oxygen species
L-NAME: NG-nitro-l-arginine methyl ester
NO: Nitric oxide
NOS: Nitric oxide synthase
OPA1: Optic Atrophy type 1
OXPHOS: Oxidative phosphorylation
PVAT: Perivascular adipose tissue
ROS: Reactive oxygen species
RAS: Renin angiotensin system
RAAS: Renin-angiotensin-aldosterone system
RT-PCR: Reverse transcription polymerase chain reaction
O_2_ ^•-^: Superoxide
SOD: Superoxide dismutase
USA: United States of America
WAT: White adipose tissue
WHO: World health organization

## Notes

### Competing Interest Statement

The authors have declared no competing interest.

